# The comprehensive detection of hemoglobinopathy variants via long-read sequencing

**DOI:** 10.1101/2024.12.03.626522

**Authors:** Jiale Xiang, Jiguang Peng, Xiangzhong Sun, Chen Jiang, Huiru Zhao, Yaya Guo, Haiyan Xu, Shanshan Gu, Haodong Ye, Long You, Xiaoyan Huang, Shiping Chen, Baosheng Zhu, Zhiyu Peng

## Abstract

**BACKGROUND:** The genetic complexity of hemoglobin genes, characterized by high GC content and homologous sequences, poses significant challenges for detecting hemoglobin variants in clinical settings.

**METHODS:** A long-read indexed PCR method utilizing the novel CycloneSEQ nanopore sequencing platform was developed to detect all variant types, including single nucleotide variants (SNVs), deletions, structural variants (SVs) in *HBA, HBB, HBD*, and *HBG* genes. The method was validated using 507 clinical samples to assess its performance.

**RESULTS:** The long-read indexed PCR system employed 13 primers targeting the hemoglobin gene clusters. This design enabled the detection of 37 types of *HBA* deletions, 5 SV (3 multicopies (αααα, ααα^*anti3.7*^, ααα^*anti4.2*^) and 2 fusion allele (*HK*αα and *anti-HK*αα)), 37 *HBB* deletions, and all SNVs in the targeted regions. Validation across 507 samples (84 with *HBA* variants, 60 with *HBB* variants, 256 with both *HBA* and *HBB* variants, and 107 with no known variants) demonstrated 100.0% sensitivity and specificity. Additionally, the long-read sequencing enabled phasing of variants within hemoglobin genes, providing insights critical for clinical interpretation.

**CONCLUSIONS:** The long-read indexed PCR method, combined with the CycloneSEQ nanopore sequencing platform, proved to be a robust and efficient solution for detecting hemoglobinopathy variants. The integration of indexed primers and barcoding enhances scalability, making this method ideal for large-scale population screening programs in the future.

## INTRODUCTION

Hemoglobinopathies are a group of inherited disorders that affect the structure or production of hemoglobin, the protein in red blood cells responsible for carrying oxygen throughout the body [1-3]. These disorders, which include conditions such as sickle cell disease and various forms of thalassemia[4], are among the most common genetic diseases worldwide [5], particularly in regions where malaria is or was endemic. The clinical manifestations of hemoglobinopathies can range from mild anemia to severe, life-threatening conditions, depending on the specific genetic mutation and its impact on hemoglobin function. Due to their high prevalence and significant health burden [6], early detection and diagnosis of hemoglobinopathies are critical for effective management and treatment, making population-based screening an essential public health strategy [7].

The molecular diagnosis of hemoglobinopathies has traditionally relied on a combination of hematological tests, such as complete blood counts and hemoglobin electrophoresis [4], along with genetic assays that target known mutations. However, these methods can be limited by their inability to detect novel or complex mutations and by the requirement for multiple tests to achieve a comprehensive diagnosis [8]. Advances in molecular techniques, such as next-generation sequencing [9, 10] and, more recently, long-read sequencing [11, 12], have significantly improved the diagnostic yield for hemoglobinopathies. These technologies allow for the simultaneous detection of a wide range of mutations, including point mutations, deletions, duplications, and other structural variants, in a single assay.

Long-read sequencing technologies, such as those developed by Pacific Biosciences [13, 14] and Oxford Nanopore Technologies [15, 16], have revolutionized genomic research by enabling the sequencing of DNA fragments that are much longer than those typically produced by traditional short-read sequencing methods [17]. Long-read sequencing offers several advantages, including the ability to capture the full sequence of the globin gene clusters, detect large deletions or insertions, and accurately characterize the diverse range of genetic mutations that underlie these disorders [11, 12]. This capability allows for more accurate assembly of complex genomic regions, identification of structural variants, and detection of phased single nucleotide polymorphisms that are difficult to resolve with short reads [18].

In the context of hemoglobinopathy screening and diagnosis, Huang et al. reported and validated the feasibility by Nanopore third-generation sequencing using 158 samples involving SNV and only 4 common α-thalassemia deletions (*--*^*THAI*^, *--*^*SEA*^, *-*α^*4.2*^, *-*α^*3.7*^), 2 β-thalassemia deletions (*SEA-HPFH*, ^*G*^γ*+(*^*A*^γδβ)^*0*^), 2 *HBA* multicopies (ααα^*anti4.2*^, ααα^*anti3.7*^), and a fusion allele (*HK*αα). However, several limitations were presented. First, some common deletions were not included in this assay including *--*^*FIL*^ on *HBA* in southern Taiwan [19], *3.5 kb deletion* on *HBB* in southern Thailand [20], *Filipino deletion* on *HBB* in Malaysia [21]. Second, samples were individually identified by barcoding sequencing in this study. Without the availability of indexed PCR primers, the scalability is limited, and the cost would be high. Moreover, phasing variants using long-read sequencing, a critical aspect for effective clinical counseling, was not addressed [11].

In this study, we developed and validated an indexed long PCR system via nanopore sequencing method for the detection of hemoglobinopathy variants. The system can be used to detect a total of 37 types α-thalassemia deletions, 37 types β-thalassemia deletions, 3 types multicopies of *HBA* (αααα, ααα^*anti3.7*^, ααα^*anti4.2*^), 2 fusion allele (*HK*αα and *anti-HK*αα) and all single nucleotide variants (SNVs) in sequencing region. Our results demonstrated the performance and stability of the method in the clinical detection of hemoglobinopathies, providing insights for the detection of hemoglobinopathies in the large-scale population.

## MATERIALS AND METHODS

### Samples and study design

The concept of the primer system was designed to cover high prevalent deletions covering hemoglobin gene cluster, and SNV variants in *HBA/HBB* genes. Moreover, we designed indexed PCR primers. Combining the barcode sequence, this system would greatly increase the scalability, which in turn decrease the cost and facilitates large-scale population screening in the future.

A total of 507 residue samples were collected, comprising 107 negative samples and 400 positive samples (carriers or patients harboring hemoglobinopathy variants). Among the positive samples, there are deletional/nondeletional (*-*α, *–*, α^*T*^α) α-thalassemia mutations, *HBA* structural variations (SVs), SNVs of *HBB* (β) and β-thalassemia deletions (*HBA*: NC_000016.10; *HBB*: NC_000011.10; reference genome: GRCh38). The experimental workflow includes multiplex long-read PCR with index primers, pooling of PCR products, ligation to barcodes and sequencing adaptors, long-read sequencing, and bioinformatic analysis (Figure 1). Long-read sequencing was performed using the CycloneSEQ WT-02, the latest single-molecule nanopore sequencer developed by BGI [22]. The study was approved by the institutional review boards of BGI (IRB24094).

**Figure 1.**
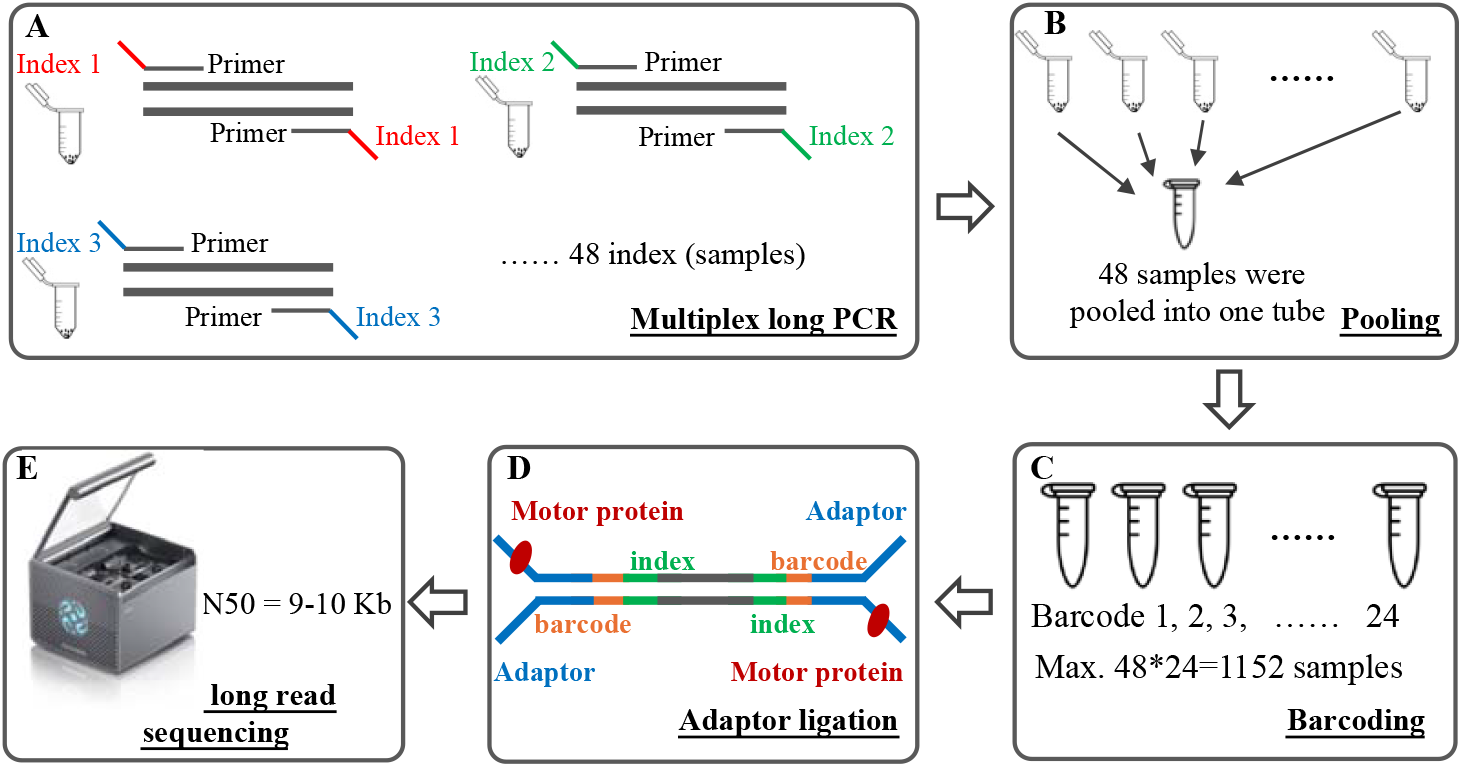
Workflow of a high-through put thalassemia screening strategy through long-read DNA sequencing. (A) Multiplex PCR of target of interest via index PCR. (B) 48 samples were pooled into a single tube. Each sample was linked with a different index. (C) The pooled product was ligated to a barcode. (D) The ligated product was then ligated to the sequencing adaptor, and then loaded onto the flow cell. (E) CycloneSEQ long-read sequencing.

### Multiplex Long-Read PCR

The optimized multiplex long-read PCR was conducted in a 25 μL reaction system, containing 20-100 ng of gDNA, 0.4 μmol/L of a mixture of PCR primers, 0.25 U of ApexHF HS DNA Polymerase CL (Accurate Biotech, Hunan, China), and 12.5 μL of 2×ApexHF CL Buffer (Accurate Biotech, Hunan, China). The PCR reaction conditions were as follows: initial denaturation at 94°C for 5 minutes, followed by 8 cycles of denaturation at 98°C for 10 seconds and extension at 68°C for 10 minutes; then 19 cycles of denaturation at 98°C for 10 seconds, annealing at 64°C for 30 seconds, annealing at 60°C for 30 seconds, and extension at 68°C for 10 minutes; and a final extension at 68°C for 10 minutes before storing the PCR products at 4°C. A mixture of 15 μL from each of the 48 multiplex long-read PCR products was purified using 0.5 volumes of VAHTS DNA Clean Beads (Vazyme Biotech, Nanjing, China) and diluted to a concentration of 40 ng/μL.

### Library Construction

Library construction was performed using the CycloneSEQ (Cyclone) kit [22]. Initially, according to the kit manual, 2 μg of purified DNA product was end-repaired using EA Enzyme (CycloneSEQ, Hangzhou, China). The product was purified using 1.0x volume of DNA clean beads and eluted with 62 μL of nuclease-free water. The concentration of the end-repaired and A-tailed product was then measured. The end-repaired and A-tailed product was then ligated with adapters using the Cyclone ligation module, purified with 1.0x volume of DNA clean beads, and eluted with 22 μL of Elution Buffer. The concentration of the ligation product was measured and used as the sequencing library.

### CycloneSEQ Long-Read Nanopore Sequencing

First, the CycloneSEQ WT-02 sequencer was started, and the chip was replaced using the chip replacement buffer, followed by chip quality control to confirm the number of active sequencing nanopores. According to the Cyclone sequencing kit manual, sequencing reagents and anchoring reagents were used to load 200 ng of the sequencing library onto the Cyclone sequencing chip. Sequencing was then initiated, and bases were called from the raw current signals, with the entire process being completed within 12-24 hours.

### Bioinformatic Analysis

First, the raw sequencing data FASTQ files generated by the CycloneSEQ-WT02 sequencer were filtered using fastp software (version 0.23.4) [23], retaining sequencing fragments with a read length greater than 1400 bp and a Phred quality score of ≥ 7. The filtered data were then subjected to the identification and splitting of sample index and amplicon primers to distinguish the sequence data of different samples and primers. For each sample data, haplotypes were pre-identified using an amplification efficiency modeling approach, then the sequencing accuracy of each haplotype’s data was corrected using an in-house bioinformatic method. Subsequently, the corrected data were aligned to the human reference genome (hg38) using Minimap2 (version 2.26-r1175) to generate BAM files, which were sorted and indexed using SAMtools (version 1.16.1) [24, 25]. Next, structural variants (αααα, ααα ^*anti3.7*^, ααα^*anti4.2*^, *HK*αα) were detected using Sniffles software (version 2.2) and CuteSV (version 2.0.3), with the results from both software tools being merged to form the final structural variant dataset [26, 27]. For complex structural variants such as alpha□globin gene triplication and Hong Kong types, the data were aligned to pre-designed multiple copy virtual genomes, allowing for accurate detection and identification of complex structural variants based on breakpoint positions. Single nucleotide variants (SNVs) were identified using freebayes software (version v1.3.6). For data that had not been pre-distinguished by haplotype, longphase software (version 1.7) was used to perform phasing in combination with SV and SNV information [28]. The phased data were then used to determine configurations and detect fusion genes. Finally, the detected variants were annotated using the thalassemia variant database(IthaNet and HbVar)[29, 30] to obtain the final detection results, and the alignment of variants to reference genome alleles was visualized using Integrative Genomics Viewer (IGV, version 2.16.2) [31].

## RESULTS

### Multiplex Long-Read PCR design

The concept of the multiplex long-read PCR for detection of hemoglobinopathy variant was shown as Figure 1-2. Specifically, 13 primers were designed to amplify variants in *hemoglobin* gene clusters (Figure 2). Detailed information of the primers was shown in Table S1. Pathogenic SNV variants in the sequencing region can be detected. Depending on the variants, the length of amplicons ranged from 1.4 kb to 16.8 kb in *HBA* including 37 types of CNVs (Table S2), and from 3.1 kb to 13.4 kb in *HBB* including 37 types of CNVs (Table S3). To validate each primer, 4 types of *HBA* CNVs (Table S2) and 12 types of *HBB* CNVs (Table S3) were selected. As shown in the Figure S1, each primer worked, and the target bands were visible via electrophoresis.

**Figure 2.**
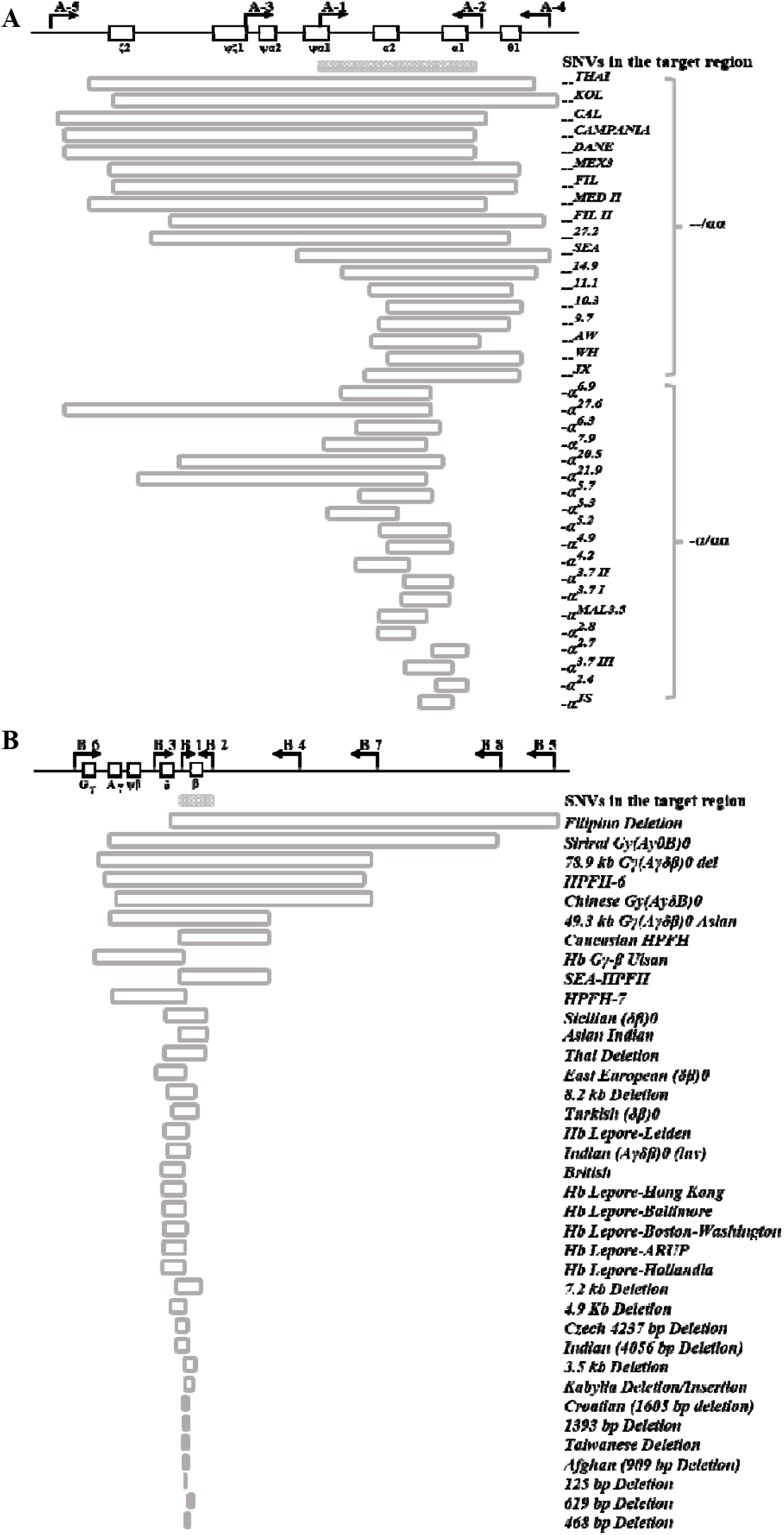
The concept of the multiplex long-read PCR for detection of variants in hemoglobin gene clusters. (A) Variants related to *HBA* and (B) *HBB*. SNV, single nucleotide variant.

### Outcomes by CycloneSEQ WT-02

CycloneSEQ WT-02 sequencer was applied for long-read sequencing of libraries of indexed long PCR products. The sequencing time was around 24 hours for 48 samples with an average data output of 10 Gb per run. The N50 of sequencing libraries was approximate 9.0 kb, which is consistent with the estimated length based on the length of the amplicons. Align the CycloneSEQ-based sequencing data with the target reference sequences of the alpha/beta-globin gene clusters, known variants were identified. The IGV plot shows the example of CNV, complex structure variants and SNVs of *HBA* and *HBB* (Figure 3). Leveraging by the length of the sequencing reads, the phase of alleles was distinguished while two variants were identified from a sample (Figure 3L).

**Figure 3.**
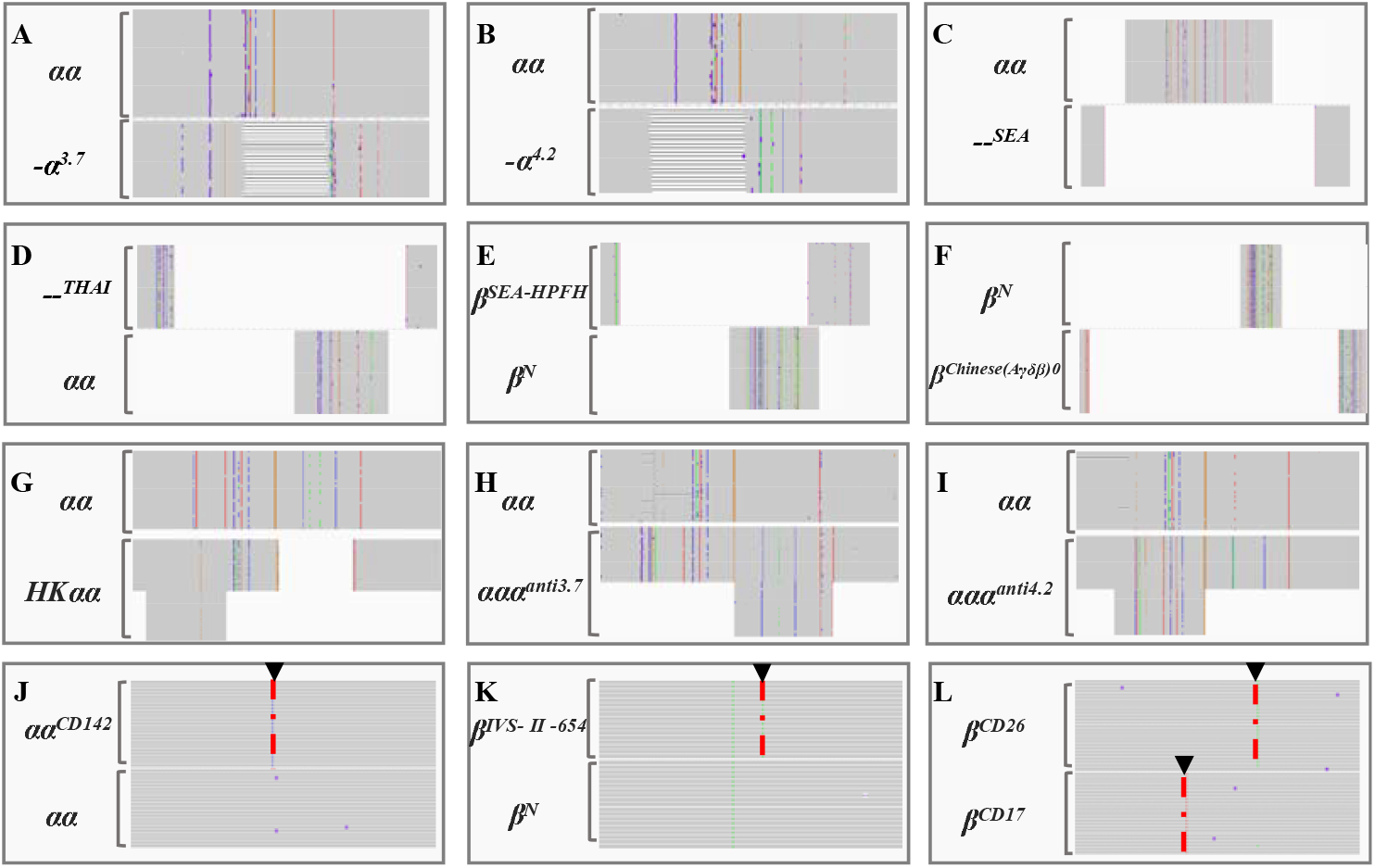
IGV plots of CNV (A-F), SV (G-I) and SNVs(J-L) with reference sequences of human genome (hg38). The normal and variant alleles are shown in each diagram. The complete gray area indicates the target sequence without deletion and variants. The blank area in the middle of the target sequence indicates a large deletion. The ▾ indicates the variation position. (A) *-*α^*3.7*^*/*αα with 3.7 kb deletion. (B) *-*α^*4.2*^*/*αα with 4.2 kb deletion. (C) *--SEA/*αα with 1.4 kb deletion. (D) -*-*^*THAI*^*/*αα with 6.0 kb deletion. (E) β^*SEA-HPFH*^*/*β^*N*^ with 7.7 kb deletion, (F) β^*Chinese (A*γδβ*)0*^*/*β^*N*^ with 8.1 kb deletion. (G) αα*/HK*αα with 4.2 kb duplication. (H) αα*/*ααα^*anti3.7*^ with 3.7 kb duplication. (I) αα*/*ααα^*anti4.2*^ with 4.2 kb duplication. (J) αα^*CD142*^ */*αα. (K) β^*IVS-II-654*^*/*β^*N*^. (L) β^*ICD26*^*/*β^*CD17*^.

### Validation of the indexed long-read sequencing method

CycloneSEQ-based long-read sequencing method not only provided high-quality data but also demonstrates stability and accuracy. Among the 507 samples with known genotypes, 107 were negative and 400 were positive (carriers of hemoglobinopathy variants). Of the 400 positive samples, 84 had *HBA* variants, 60 had *HBB* variants, and 256 had both *HBA* and *HBB* variants (Table 1). From the perspective of variant types, the 400 positive samples had 60 types of variants including 9 *HBA* CNV (*--*^*FIL*^,*--*^*SEA*^, *--*^*THAI*^, *-*α^*3.7□*^, *-*α^*4.2*^, *-*α^*21.9*^, *-*α^*27.6*^, *-*α^*27.2*^), 3 *HBA* multicopies (ααα^*anti3.7*^, ααα^*anti4.2*^, αααα), 1 *HBA* fusion allele (*HK*αα), 16 *HBA* SNV, 4 *HBB* CNV (*3.5 kb deletion, Chinese (A*γδβ*)0, SEA-HPFH*, and *Taiwanese deletion*) and 27 *HBB* SNV (Table 2 and Table S3). The concordance rate of the long-read sequencing method was 100.0% compared to the known results. Overall, the sensitivity and specifically was 100.0%.

**Table 1.**
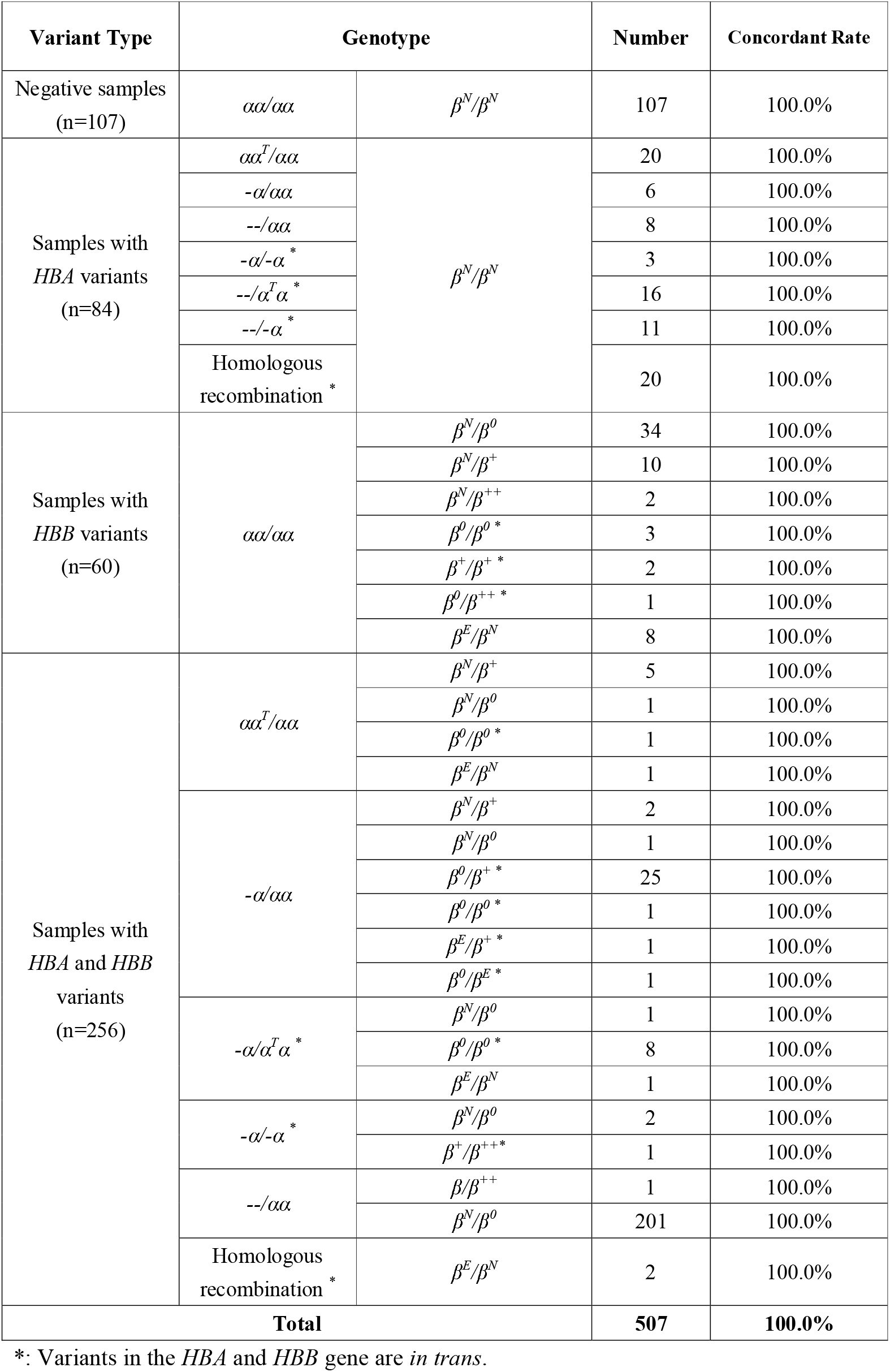
The performance of the long-read sequencing method in the detection of α- and β-thalassemia variants by CycloneSEQ.

| Variant Type | Genotype |  | Number | Concordant Rate |
| --- | --- | --- | --- | --- |
| Negative samples<br>(n=107) | $\alpha\alpha/\alpha\alpha$ | $\beta^N/\beta^N$ | 107 | 100.0% |
| Samples with<br>*HBA* variants<br>(n=84) | $\alpha\alpha^T/\alpha\alpha$ | $\beta^N/\beta^N$ | 20 | 100.0% |
| | $-\alpha/\alpha\alpha$ | | 6 | 100.0% |
| | $--/\alpha\alpha$ | | 8 | 100.0% |
| | $-\alpha/-\alpha^*$ | | 3 | 100.0% |
| | $--/\alpha^T\alpha^*$ | | 16 | 100.0% |
| | $--/-\alpha^*$ | | 11 | 100.0% |
|  | Homologous<br>recombination * |  | 20 | 100.0% |
| Samples with<br>*HBB* variants<br>(n=60) | $\alpha\alpha/\alpha\alpha$ | $\beta^N/\beta^0$ | 34 | 100.0% |
| | | $\beta^N/\beta^+$ | 10 | 100.0% |
| | | $\beta^N/\beta^{++}$ | 2 | 100.0% |
| | | $\beta^0/\beta^{0*}$ | 3 | 100.0% |
| | | $\beta^+/\beta^{+*}$ | 2 | 100.0% |
| | | $\beta^0/\beta^{++*}$ | 1 | 100.0% |
| | | $\beta^E/\beta^N$ | 8 | 100.0% |
| Samples with<br>*HBA* and *HBB*<br>variants<br>(n=256) | $\alpha\alpha^T/\alpha\alpha$ | $\beta^N/\beta^+$ | 5 | 100.0% |
| | | $\beta^N/\beta^0$ | 1 | 100.0% |
| | | $\beta^0/\beta^{0*}$ | 1 | 100.0% |
| | | $\beta^E/\beta^N$ | 1 | 100.0% |
| | $-\alpha/\alpha\alpha$ | $\beta^N/\beta^+$ | 2 | 100.0% |
| | | $\beta^N/\beta^0$ | 1 | 100.0% |
| | | $\beta^0/\beta^{+*}$ | 25 | 100.0% |
| | | $\beta^0/\beta^{0*}$ | 1 | 100.0% |
| | | $\beta^E/\beta^{+*}$ | 1 | 100.0% |
| | | $\beta^0/\beta^E$ | 1 | 100.0% |
| | $-\alpha/\alpha^T\alpha^*$ | $\beta^N/\beta^0$ | 1 | 100.0% |
| | | $\beta^0/\beta^{0*}$ | 8 | 100.0% |
| | | $\beta^E/\beta^N$ | 1 | 100.0% |
| | $-\alpha/-\alpha^*$ | $\beta^N/\beta^0$ | 2 | 100.0% |
| | | $\beta^+/\beta^{++}$ | 1 | 100.0% |
| | $--/\alpha\alpha$ | $\beta/\beta^{++}$ | 1 | 100.0% |
| | | $\beta^N/\beta^0$ | 201 | 100.0% |
| | Homologous<br>recombination * | $\beta^E/\beta^N$ | 2 | 100.0% |
| **Total** |  |  | **507** | **100.0%** |

\*: Variants in the *HBA* and *HBB* gene are *in trans*.

**Table 2.**
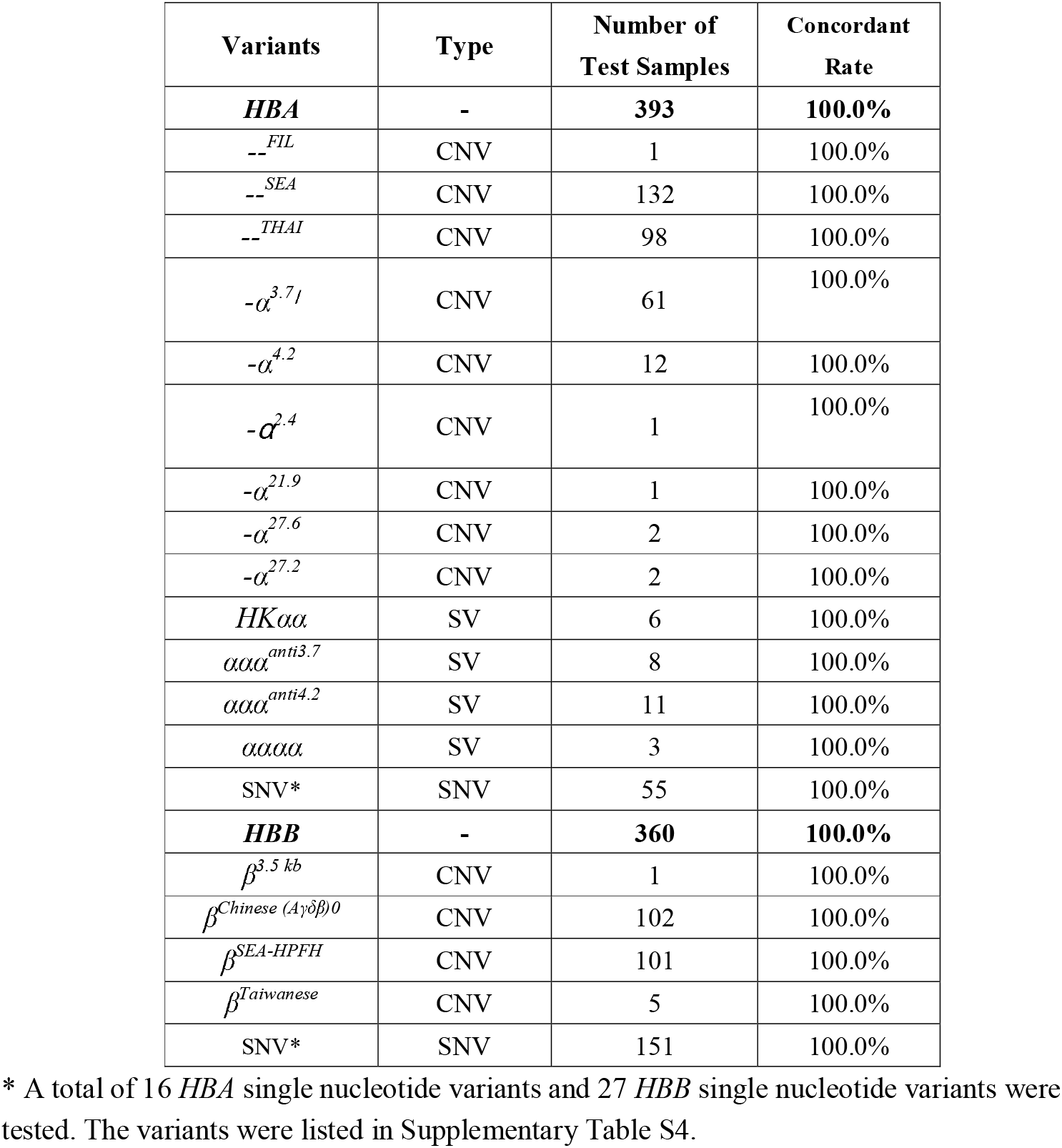
Variant types of tested samples.

## DISCUSSION

In this study, we developed a novel method based on the new nanopore sequencing platform, CycloneSEQ [22], for the detection of hemoglobinopathy variants. A key advantage of this method is the development of indexed primers for each amplicon (n=48), allowing the index sequences to differentiate samples separately. With the availability of sequencing barcodes (n=24), which is generally ligated to indexed PCR products, this design significantly increases the capacity for sample testing, allowing the feasibility of large-scale population screening (max. 48*24=1152 samples) in the future. To our knowledge, this is the first study reported the indexed long-read amplicon sequencing for monogenic disorders.

To validate the method, we analyzed 507 clinical samples encompassing 60 variant types, including 29 variants in *HBA* and 31 variants in *HBB* (detailed in Table 2 and Table S4). These variants spanned multiple types, such as CNVs, SNVs, and SVs, covering nearly all common hemoglobinopathy-related variants. To the best of our knowledge, this study represents the most extensive validation effort to date for detecting hemoglobinopathy variants via long-read sequencing. Our findings confirm that the long-range indexed PCR system is a robust and practical solution for this application.

Leveraging the flexibility of nanopore sequencing, each flow cell can process 12-48 samples per run. The total turnaround time is approximately 33 hours for 48 samples, 21 hours for 24 samples, and 15 hours for 12 samples, encompassing DNA extraction and multiplex long-range indexed PCR (6 hours), library preparation (2.5 hours), and sequencing with data analysis (about 24 hours for 48 samples, 12 hours for 24 samples, and 6 hours for 12 samples). Between runs, each flow cell was washed using the Flow Cell Wash Kit to restore sequencing pore activity. Contamination risks between sequential runs were minimized through sample indexing, allowing effective filtering of residual sequences from previous runs.

One notable advantage of long-read sequencing is its ability to determine the configuration of two variants identified within the same gene [32]. Traditional phasing methods using short-read sequencing heavily rely on paternal and maternal samples. However, phasing variants *in trans* or *cis* configurations is critical for understanding disease mechanisms and guiding clinicians in providing precise counseling [33]. In this study, 53 samples harbored two *HBA* variants (26 CNV+SNV, 16 CNV+CNV, 4 SV+CNV, 1 SV+SNV, 6 SV+SV), while 44 samples harbored two *HBB* variants (1 CNV+SNV, 43 SNV+SNV). Using long-read sequencing data, we successfully resolved the *trans* configuration for all tested samples.

In conclusion, the long-read indexed PCR method using the novel CycloneSEQ nanopore sequencing platform proved to be a reliable and efficient approach for detecting hemoglobinopathy variants. Thanks to its ability to generate long reads, this method can accurately identify all variant types, including SNVs, CNVs, SVs, and fusion alleles, while also determining the phase of variants. Furthermore, the integration of indexed primers with barcodes enhances the scalability of the assay, making it well-suited for large-scale population screening programs in the future.

## Supporting information

Supplementary materials provide additional support and context to the main text of the article.

