## Supplementary materials provide additional support and context to the main text of the article. for "The comprehensive detection of hemoglobinopathy variants via long-read sequencing"

**Table S1 Primers and amplicons of multiplex long PCR**

| **Primer name** | **Sequence (5‘-3’)** | **Position (hg38)** |
| --- | --- | --- |
| A-1 | GTTTACCCATGTGGTGCCTCCAT | Chr16: 169269-169291 |
| A-2 | GCATGAGTCATCACACCTGGACAAT | Chr16: 178806-178830 |
| A-3 | GAGCGATCTGGGCTCTGTGTTCTC | Chr16: 165245-165277 |
| A-4 | GCACATGTGCTTTCCTCTTCGTCTTC | Chr16: 185947-185972 |
| A-5 | GATCTGAGGTGGCACACAAGCAT | Chr16: 146487-146509 |
| B-1 | CTGCGTGTGTGAGAATAATCAGAGT | Chr11: 5228685-5228709 |
| B-2 | CTTTGTAGGTCACTCAGGACTTGTT | Chr11: 5237748-5237772 |
| B-3 | TAGTTTGAACTCACCTCTGGTTACT | Chr11: 5236060-5236084 |
| B-4 | TGCTAGGGCTTACTGAAGTAATCAT | Chr11: 5200658-5200682 |
| B-5 | TTGTCTATTGCAGGTGTGTAGACAT | Chr11: 5112671-5112695 |
| B-6 | ACAGTCCCTGCCTCTTAAGAGTT | Chr11: 5255756-5255778 |
| B-7 | GCTCTTGGGTGTACAAGTTTTCCAT | Chr11: 5168742-5168766 |
| B-8 | AGAATTCTCAGGCATTGTCCACTGT | Chr11: 5131111-5131135 |

Chr, chromosome

**Table S2 Summary of *HBA* variants and length of amplicons.**

| **#** | **Gene** | **Common CNV/SV Name** | **Forword Primer & Reverse Primer** | **Length of Amplicon（kb）** | **Comment** |
| --- | --- | --- | --- | --- | --- |
| 1 | *HBA* | *αα* | A-1&A-2 | 9.6 | Used to validate the effectiveness of the primers |
| 2 | *HBA* | *-α^3.7I^* | A-1&A-2 | 5.8 |  |
| 3 | *HBA* | *--^SEA^* | A-3&A-4 | 1.4 |  |
| 4 | *HBA* | *--^THAI^* | A-5&A-4 | 6.0 |  |
| 5 | *HBA* | *-α^3.7 Ⅱ^* | A-1&A-2 | 5.8 | / |
| 6 | *HBA* | *-α^3.7 Ⅲ^* | A-1&A-2 | 5.8 |  |
| 7 | *HBA* | *--^JX^* | A-1&A-4 | 3.7 |  |
| 8 | *HBA* | *-α^11.1^* | A-1&A-4 | 5.5 |  |
| 9 | *HBA* | *-α^10.3^* | A-1&A-4 | 6.4 |  |
| 10 | *HBA* | *-α^9.7^* | A-1&A-4 | 7.0 |  |
| 11 | *HBA* | *--^AW^* | A-1&A-4 | 8.5 |  |
| 12 | *HBA* | *-α^5.7^* | A-1&A-2 | 3.9 |  |
| 13 | *HBA* | *-α^JS^* | A-1&A-2 | 7.0 |  |
| 14 | *HBA* | *-α^MAL3.5^* | A-1&A-2 | 6.0 |  |
| 15 | *HBA* | *-α^2.8^* | A-1&A-2 | 6.7 |  |
| 16 | *HBA* | *--^14.9^* | A-3&A-4 | 5.8 |  |
| 17 | *HBA* | *--^FIL^* | A-5&A-4 | 8.8 |  |
| 18 | *HBA* | *--^MED II^* | A-5&A-4 | 9.5 |  |
| 19 | *HBA* | *--^FIL II^* | A-5&A-4 | 10.4 |  |
| 20 | *HBA* | *--^27.2^* | A-5&A-4 | 12.2 |  |
| 21 | *HBA* | *-α^27.6^* | A-5&A-2 | 4.7 |  |
| 22 | *HBA* | *-α^21.9^* | A-5&A-2 | 10.4 |  |
| 23 | *HBA* | *-α^20.5^* | A-5&A-2 | 16.8 |  |
| 24 | *HBA* | *-α^7.9^* | A-3&A-2 | 5.6 |  |
| 25 | *HBA* | *-α^6.9^* | A-3&A-2 | 6.7 |  |
| 26 | *HBA* | *--^KOL^* | A-5&A-4 | 6.1 |  |
| 27 | *HBA* | *--^CAL^* | A-5&A-4 | 7.0 |  |
| 28 | *HBA* | *--^CAMPANIA^* | A-5&A-4 | 8.6 |  |
| 29 | *HBA* | *--^MEX3^* | A-5&A-4 | 8.7 |  |
| 30 | *HBA* | *-α^2.7^* | A-1&A-2 | 6.9 |  |
| 31 | *HBA* | *-α^6.3^* | A-1&A-2 | 3.2 |  |
| 32 | *HBA* | *-α^5.3^* | A-3&A-2 | 8.3 |  |
| 33 | *HBA* | *-α^5.2^* | A-1&A-2 | 4.4 |  |
| 34 | *HBA* | *-α^4.9^* | A-1&A-2 | 4.7 |  |
| 35 | *HBA* | *-α^4.2^* | A-1&A-2 | 5.3 |  |
| 36 | *HBA* | *-α^2.4^* | A-1&A-2 | 7.2 |  |
| 37 | *HBA* | *--^DANE^* | A-5&A-4 | 8.6 |  |
| 38 | *HBA* | *--^WH^* | A-1& A-4 | 6.5 |  |
| 39 | *HBA* | *ααα^anti3.7^* | A-1&A-2 | 13.3 |  |
| 40 | *HBA* | *ααα^anti4.2^* | A-1&A-2 | 13.8 |  |
| 41 | *HBA* | *HKαα* | A-1&A-2 | 10.0 |  |
| 42 | *HBA* | *αααα* | A-1&A-2 | 18.0 |  |
| 43 | *HBA* | *Anti-HKαα* | A-1&A-2 | 9.3 |  |

CNV, copy number variation;

SV, structure variation.

**Table S3 Summary of *HBB* variants and length of amplicons.**

| **#** | **Gene** | **Common CNV/SV Name** | **Forword Primer & Reverse Primer** | **Length of Amplicon（kb）** | **Comment** |
| --- | --- | --- | --- | --- | --- |
| 1 | *HBB* | *βN* | B-1&B-2 | 9.1 | Used to validate the effectiveness of the primers |
| 2 | *HBB* | *Taiwanese deletion* | B-1&B-2 | 6.1 |  |
| 3 | *HBB* | *3.5 kb deletion* | B-1& B-2 | 5.6 |  |
| 4 | *HBB* | *Hb Lepore-Leiden* | B-3&B-2 | 8.9 |  |
| 5 | *HBB* | *Thai deletion* | B-3&B-2 | 3.8 |  |
| 6 | *HBB* | *Sicilian (δβ)0* | B-3&B-2 | 3.1 |  |
| 7 | *HBB* | *Turkish (δβ)0* | B-3&B-2 | 8.8 |  |
| 8 | *HBB* | *Chinese Gy(AyδB)0* | B-6&B-7 | 8.2 |  |
| 9 | *HBB* | *HPFH-6* | B-6&B-7 | 7.8 |  |
| 10 | *HBB* | *SEA-HPFH* | B-3&B-4 | 8.0 |  |
| 11 | *HBB* | *Filipino deletion* | B-3&B-5 | 4.9 |  |
| 12 | *HBB* | *Sirirai Gy(AyôB)0* | B-6&B-8 | 6.7 |  |
| 13 | *HBB* | *Indian (Aγδβ)0 (inv)* | B-3&B-2 | 9.0 |  |
| 14 | *HBB* | *78.9 kb Gγ(Aγδβ)0 del* | B-6&B-7 | 3.2 | / |
| 15 | *HBB* | *49.3 kb Gγ(Aγδβ)0 Asian del* | B-6&B-4 | 5.8 |  |
| 16 | *HBB* | *Caucasian HPFH* | B-3&B-4 | 7.6 |  |
| 17 | *HBB* | *Hb Gγ-β Ulsan* | B-6&B-2 | 8.4 |  |
| 18 | *HBB* | *HPFH-7(Hb Kenya)* | B-6&B-2 | 13.4 |  |
| 19 | *HBB* | *East European (δβ)0* | B-3&B-2 | 7.3 |  |
| 20 | *HBB* | *8.2 kb deletion* | B-3&B-2 | 8.2 |  |
| 21 | *HBB* | *Asian Indian* | B-3&B-2 | 6.1 |  |
| 22 | *HBB* | *British* | B-3&B-2 | 9.0 |  |
| 23 | *HBB* | *Hb Lepore-Hong Kong* | B-3&B-2 | 9.1 |  |
| 24 | *HBB* | *Hb Lepore-Baltimore* | B-3&B-2 | 9.0 |  |
| 25 | *HBB* | *Hb Lepore-Boston-Washington* | B-3&B-2 | 9.0 |  |
| 26 | *HBB* | *Hb Lepore-ARUP* | B-3&B-2 | 9.0 |  |
| 27 | *HBB* | *Hb Lepore-Hollandia* | B-3&B-2 | 9.0 |  |
| 28 | *HBB* | *7.2 kb deletion* | B-3&B-2 | 9.2 |  |
| 29 | *HBB* | *4.9 Kb deletion* | B-3&B-2 | 11.6 |  |
| 30 | *HBB* | *Czech 4237 bp deletion* | B-3&B-2 | 12.2 |  |
| 31 | *HBB* | *Indian (4056 bp deletion)* | B-3&B-2 | 12.4 |  |
| 32 | *HBB* | *Kabylia deletion/insertion* | B-1&B-2 | 7.0 |  |
| 33 | *HBB* | *Croatian (1605 bp deletion)* | B-1&B-2 | 7.5 |  |
| 34 | *HBB* | *1393 bp deletion* | B-1&B-2 | 7.7 |  |
| 35 | *HBB* | *Afghan (909 bp deletion)* | B-1&B-2 | 8.2 |  |
| 36 | *HBB* | *125 bp deletion* | B-1&B-2 | 9.0 |  |
| 37 | *HBB* | *468 bp deletion* | B-1&B-2 | 8.7 |  |
| 38 | *HBB* | *619 bp deletion* | B-1&B-2 | 8.5 |  |

**Table S4 Single nucleotide variants (SNV) of tested samples**

| **SNV** | **Number of Test** | **Concordant Rate** |
| --- | --- | --- |
| **HBA** | **55** | **100.0%** |
| *HBA1:c.334G>T* | 1 | 100.0% |
| *HBA1:c.364G>A* | 10 | 100.0% |
| *HBA2:c.369C>G* | 12 | 100.0% |
| *HBA2:c.377T>C* | 4 | 100.0% |
| *HBA2:c.40G>T* | 1 | 100.0% |
| *HBA2:c.427T>C* | 11 | 100.0% |
| *HBA1:c.46G>C* | 2 | 100.0% |
| *HBA1:c.51G>C* | 1 | 100.0% |
| *HBA2:c.54delC* | 1 | 100.0% |
| *HBA1:c.84G>T* | 5 | 100.0% |
| *HBA2:c.95G>A* | 2 | 100.0% |
| *HBA1:c.178G>A* | 1 | 100.0% |
| *HBA1:c.223G>C* | 1 | 100.0% |
| *HBA2:c.275T>C* | 1 | **100.0%** |
| *HBA2:c.80C>A* | 1 | 100.0% |
| *HBA1:c.104T>G* | 1 | 100.0% |
| **HBB** | **151** | **100.0%** |
| *HBB:c.-79A>G* | 1 | 100.0% |
| *HBB:c.-81A>C* | 1 | 100.0% |
| *HBB:c.-122T>A* | 1 | 100.0% |
| *HBB:c.-136C>G* | 1 | 100.0% |
| *HBB:c.-78A>G* | 7 | 100.0% |
| *HBB:c.315+5G>C* | 1 | 100.0% |
| *HBB:c.315+1G>T* | 1 | 100.0% |
| *HBB:c.315+2del* | 1 | 100.0% |
| *HBB:c.316-197C>T* | 36 | 100.0% |
| *HBB:c.92+1G>T* | 3 | 100.0% |
| *HBB:c.*129T>A* | 5 | 100.0% |
| *HBB:c.48G>A* | 1 | 100.0% |
| *HBB:c.113G>A* | 1 | 100.0% |
| *HBB:c.165_177delTATGGGCAACCCT* | 1 | 100.0% |
| *HBB:c.208G>A* | 2 | 100.0% |
| *HBB:c.68A>C* | 1 | 100.0% |
| *HBB:c.188C>T* | 1 | 100.0% |
| *HBB:c.199A>G* | 1 | 100.0% |
| *HBB:c.170G>A* | 2 | 100.0% |
| *HBB:c.343C>A* | 1 | 100.0% |
| *HBB:c.85dupC* | 5 | 100.0% |
| *HBB:c.394C>A* | 1 | 100.0% |
| *HBB:c.217dupA* | 27 | 100.0% |
| *HBB:c.79G>A* | 5 | 100.0% |
| *HBB:c.126_129delCTTT* | 18 | 100.0% |
| *HBB:c.90C>T* | 8 | 100.0% |
| *HBB:c.52A>T* | 18 | 100.0% |

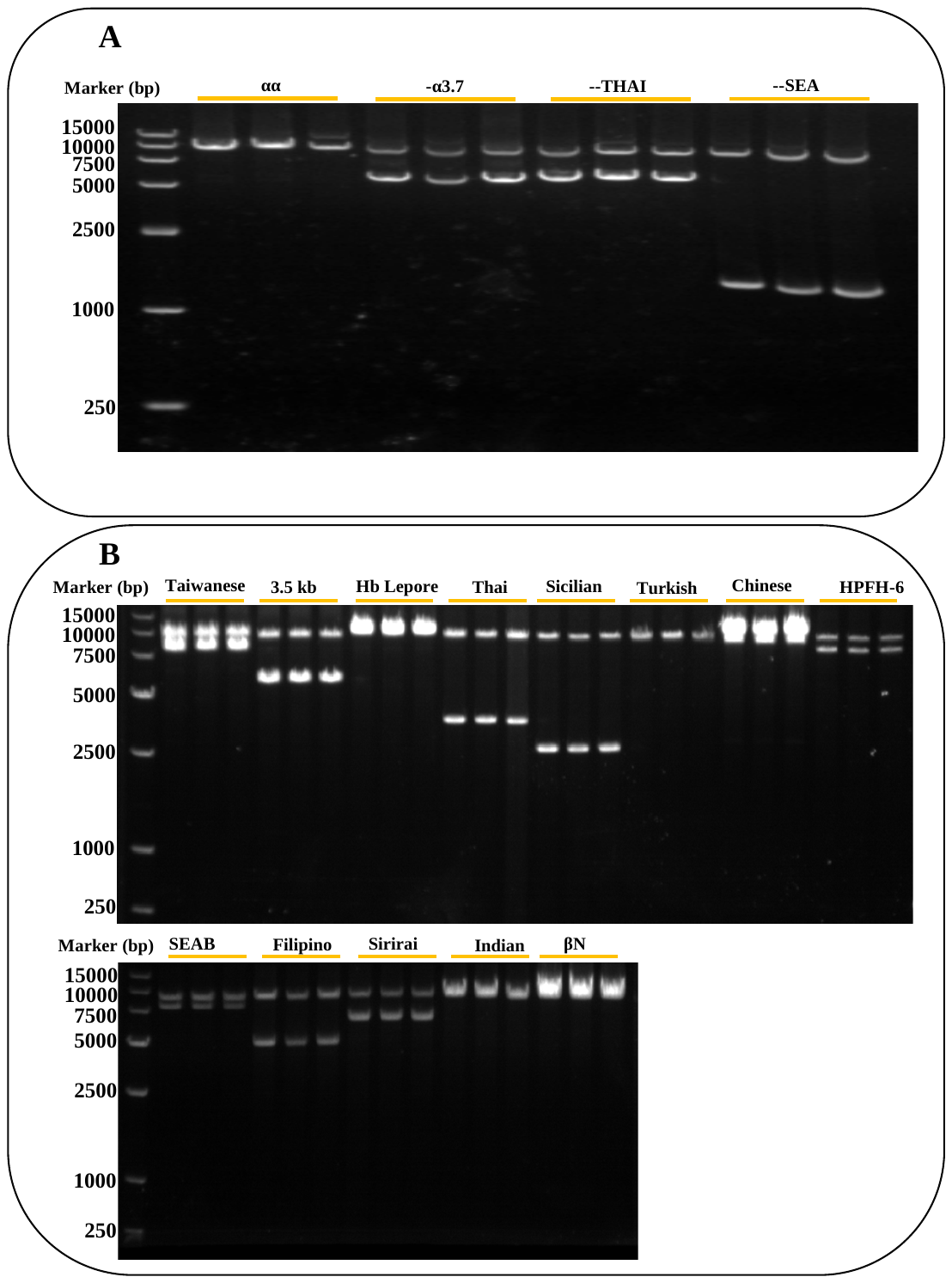

**Figure S1 Electrophoresis results of target region in 0.8% agarose gel.**
